# Rapid evolution of the inter-sexual genetic correlation for fitness in *Drosophila melanogaster*

**DOI:** 10.1101/032664

**Authors:** J. M. Collet, S. Fuentes, J. Hesketh, M. S. Hill, P. Innocenti, E. H. Morrow, K. Fowler, M. Reuter

## Abstract

Sexual antagonism (SA) arises when male and female phenotypes are under opposing selection, yet genetically correlated. Until resolved, antagonism limits evolution towards optimal sex-specific phenotypes. Despite its importance for sex-specific adaptation and existing theory, the dynamics of SA resolution are not well understood empirically. Here, we present data from *Drosophila melanogaster*, compatible with a resolution of SA. We compared two independent replicates of the ‘LH_M_’ population in which SA had previously been described. Both had been maintained under identical, controlled conditions, and separated for <250 generations. Although heritabilities of male and female fitness were similar, the inter-sexual genetic correlation differed significantly, being negative in one replicate (indicating SA) but close to zero in the other. Using population sequencing, we show that phenotypic differences were associated with population divergence in allele frequencies at non-random loci across the genome. Large frequency changes were more prevalent in the population without SA and were enriched at loci mapping to genes previously shown to have a sexually antagonistic relationships between expression and fitness. Our data suggest that rapid evolution towards SA resolution has occurred in one of the populations and open avenues towards studying the genetics of SA and its resolution.

## Introduction

Due to their different reproductive roles, male and female adults are often selected for different optimal phenotypes. However, the response to this divergent selection is limited by the fact that the sexes share a large part of their genome and, thus, new mutations frequently affect the phenotype of males and females in a similar way. The combination of genetically correlated male and female phenotypes and divergent selection on the sexes sets the scene for intra-locus sexual conflict or sexual antagonism (SA), where some alleles increase the fitness in one sex at the expense of the fitness in the other sex (Rice 1984; Bonduriansky and Chenoweth 2009; Van Doorn 2009; Connallon and Clark 2014). Sexually antagonistic genetic variation has been shown to segregate in natural and laboratory populations of a wide range of organisms, including insects (Chippindale et al. 2001; Fedorka and Mousseau 2004; Lewis et al. 2011; Berg and Maklakov 2012), vertebrates (Brommer et al. 2007; Foerster et al. 2007; Mainguy et al. 2009; Mokkonen et al. 2011) and plants (Kohorn 1994; Scotti and Delph 2006; Delph et al. 2011). This growing body of evidence demonstrates that the common genetic basis of male and female phenotypes limits the adaptive evolution of sex-specific traits. The adaptive trade-offs inherent in sexually antagonistic allelic variation prevents both sexes from attaining their sex-specific optima and generates balancing selection that can maintain genetic variation at antagonistic loci. For this reason, sexual antagonism is also a powerful agent for the maintenance of genetic variation for fitness (Kidwell et al. 1977).

Despite being recognised as an important evolutionary force, the study of SA has so far focused mostly on characterising antagonism as snapshots of particular populations at particular time-points and relatively little is known about its long-term dynamics (Stewart et al. 2010; Dean et al. 2012; Pennell and Morrow 2013). More specifically, it is currently unclear to what degree the intensity of antagonism changes over time, and over what timescale such changes occur. Similarly, there is scant information on whether individual antagonistic loci remain polymorphic over long periods of time and, if not, what mechanisms are involved in the fixation of one or the other allele. Answering these questions is vital for our understanding of ongoing conflicts over adaptation between the sexes, and of the evolution of sexual dimorphism.

Changes in the extent of SA can occur in response to variation in a population’s environment as well as in its genetic composition. Experimental work in fruitflies has shown that the extent of SA varies between environmental conditions due to genotype-by-environment interactions (Delcourt et al. 2009; Punzalan et al. 2014). Large shifts in the environment might also eliminate antagonism (Long et al. 2012; Connallon and Clark 2014; Punzalan et al. 2014). This occurs if the change in conditions is large enough for the new selective optima of two sexes to both fall above (or below) their current trait values. This would then put both sexes under concordant directional selection and result in a positive genetic correlation between male and female fitness across genotypes.

The extent of SA can also vary in response to changes in the genetic composition of a population through mutation, drift and selection. Mutations can increase the degree of antagonism if it generates similar phenotypic effects in the two sexes and hence increases their genetic correlation for traits under opposing selection (Connallon and Clark 2014). Genetic drift, in contrast, will result in a loss of genetic polymorphism at antagonistic loci, and hence can reduce the degree of SA observable within populations (Connallon and Clark 2012; Mullon et al. 2012; Hesketh et al. 2013). The rate at which this loss occurs depends on the effective population size at antagonistic loci, and thus will be faster in small populations, populations with large reproductive skew, and at loci under increased drift due to effects of chromosome dose (such as X or Z sex chromosomes) or selective interference between loci (Vicoso and Charlesworth 2009; Connallon and Clark 2012; Mullon et al. 2012).

Finally, and most importantly from an evolutionary point of view, the intensity of SA can be reduced through adaptive evolution. The trade-off between male and female fitness that underlies antagonism will create a selection pressure for mechanisms that diminish the deleterious fitness effects of the allele whenever it resides in the disfavoured sex. These mechanisms are not well characterised empirically but could include sex-specific modifier loci (Rice 1984), sex-specific dominance (Barson et al. 2015), gene duplication followed by the evolution of sex-specific gene expression and adaptation in the two paralogues (Connallon and Clark 2011; Parsch and Ellegren 2013), genetic imprinting (Iwasa and Pomiankowski 1999, 2001; Day and Bonduriansky 2004) or sex-specific splicing (Pennell and Morrow 2013). Such adaptations will result in the long-term resolution of antagonism and allow both sexes to approach their phenotypic optima, thereby increasing the degree of sexual dimorphism (Lande 1980; Ellegren and Parsch 2007; Bonduriansky and Chenoweth 2009; Parsch and Ellegren 2013).

Understanding how antagonism evolves requires repeated measurements of SA at several time points. This can be achieved by monitoring SA in a single population through time or, alternatively, by measuring the extent of SA in different populations derived from a common ancestral population. Here, we present the results of such a comparative study investigating SA in two recently diverged replicate populations of the laboratory-adapted *D. melanogaster* stock LH_M_. Both replicates were derived from the original LH_M_ population maintained by William Rice at the University of California, Santa Barbara, in which sexual antagonism had previously been documented (Chippindale et al. 2001). At the time of our analysis, they had been separated for about 200 generations but maintained according to the same strictly imposed rearing regime (see Methods, Fig. 1). Combining existing data on the genetic architecture of male and female fitness in one population (LH_M_-UU, Innocenti and Morrow 2010) with newly collected data on the other (LH_M_-UCL), we show that the inter-sexual genetic correlation for fitness has diverged significantly between the two populations since they have been separated. While one population shows a negative genetic correlation between male and female fitness, indicative of antagonism, male and female fitness are not significantly correlated in the other, despite similar levels of heritable variation for fitness in the two sexes of both populations. Using a population genomic approach to compare allele frequencies at genome-wide SNP loci, we identify islands of significant differentiation between the populations, both on the X chromosome and the autosomes. We show that patterns of differentiation are biased towards frequency change in the population with reduced SA and enriched in genes with sexually antagonistic expression patterns. We argue that these results are unlikely to be due to environmental effects and compatible with the alleviation or resolution of sexual antagonism in one of the study populations. Given the increasing recognition of SA as a significant evolutionary force, our findings provide insights into the long-term evolutionary dynamics of SA and indicate future ways to elucidate the mechanistic basis of antagonism.

**Figure 1.**
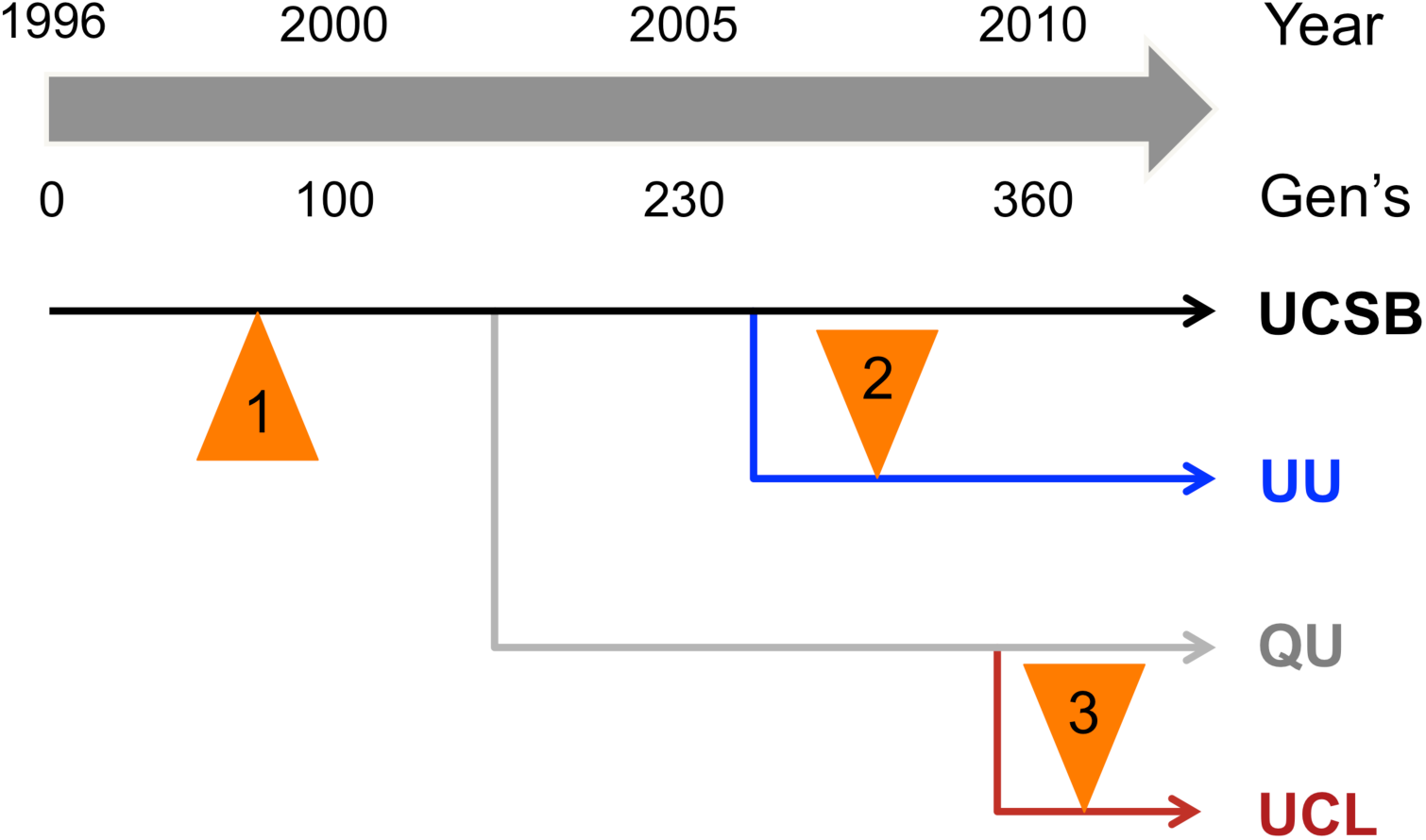
Schematic history of LH_M_ populations. The relationship between the LH_M_ populations used here, and the ancestral populations from which they were derived. The timeline is represented in calendar years and generations (approximate). For more details, refer to the Methods section of the main text. Orange triangles denote quantitative genetic studies of sex-‐specific fitness, 1: Chippindale et al. (2001), 2: Innocenti and Morrow (2010), 3: this study.

## Materials and Methods

### Study populations

This study uses two replicate populations derived from the outbred laboratory stock LH_M_ (here LH_M_-UCSB), maintained by W. Rice at the University of California, Santa Barbara (Fig. 1). LH_M_-UCL was established from a duplicate of LH_M_-UCSB that was taken to Queen’s University by A. Chippindale in February 2002 and then transferred to the Reuter group in May 2009. Independently, a replicate of LH_M_-UCSB was taken to the Morrow group, University of Uppsala, in December 2005 to establish the other population, LH_M_-UU.

Starting with the establishment of LH_M_-UCSB in 1996, all LH_M_ populations have been continuously maintained at a constant adult population size of 1792 individuals (896 males and 896 females) and under the same strictly regimented 14-day rearing regime (described in Rice et al. 2005). Therefore, neither LH_M_-UCL nor LH_m_-UU had experienced population bottlenecks or more than subtle environmental shifts. The rigorous two-week cycle further allowed us to estimate the number of generations of separation between the populations, dating back to the split between LH_M_-QU and in February 2002 (Fig. 1). At the time of sampling genotypes for fitness measurements in October 2007, LH_M_-UU had undergone approximately 145 generations from that branching point. In LH_M_-UCL, genotypes were sampled in June 2010, approximately 215 generations since the original split.

### Hemiclonal analysis of male and female fitness

We used hemiclonal analysis to measure the effects of haploid genomes on male and female fitness (see Abbott and Morrow 2011 for a review of the approach). Hemiclonal individuals share a common copy of chromosomes X, 2 and 3, which amount to 99.5% of an identical haplotype (all genes except for the 0.5% of the genome located on the ‘dot’ fourth chromosome). The quantitative genetic analysis captures additive genetic effects of the hemiclonal X, 2 and 3 chromosomes (as well as additive effects of epistatic interactions between alleles within the hemiclonal haplotype) but averages the effects of epistasis or dominance between the hemiclonal haplotype and the genetic background (Rice et al. 2005).

The quantitative genetic analysis of fitness used here was closely modelled on previous studies (Chippindale et al. 2001; Pischedda and Chippindale 2006; Innocenti and Morrow 2010; see Supplementary Material and the previously cited studies for details). To measure fitness in hemiclones, crosses were performed to generate males and females that carry an identical hemiclonal genome complemented with random genetic material from the corresponding source population. The fitness of these flies was then measured under conditions that closely mimic the LH_M_ rearing regime and in competition with a standard competitor stock. The competitors provide a point of reference from which to calculate the relative fitness of different experimental genotypes. Their exact identity and genetic composition is not important for the results generated, as long as their fitness is similar to those of the experimental flies. The competitor flies used here carried a homozygous *brown (bw)* mutation in a variable, outbred LH_M_ background, ensuring competitiveness while allowing us to assign paternity experimental and competitor males. The competitior stock was maintained following exactly the same regime as the wildtype LH_M_ population.

**Fitness measurements for the LH_M_-UCL population:** LH_M_-UCL hemiclonal lines were established in June 2010 and their fitness was measured between July 2010 and September 2011. For all lines, fitness was assayed three times in each sex in blocks that included one replicate of each hemiclonal line and alternatingly assayed male and female fitness. Complete fitness data was obtained for 113 lines. In order to assess potential genotype-by-laboratory effects, we also assessed the fitness of nine of the ten most sexually antagonistic lines created in the LH_M_-UU population (Fig. 1 in Innocenti and Morrow 2010; see also section ‘Comparing fitness measures across laboratories’ below and Figs. 2 and S1).

**Figure 2.**
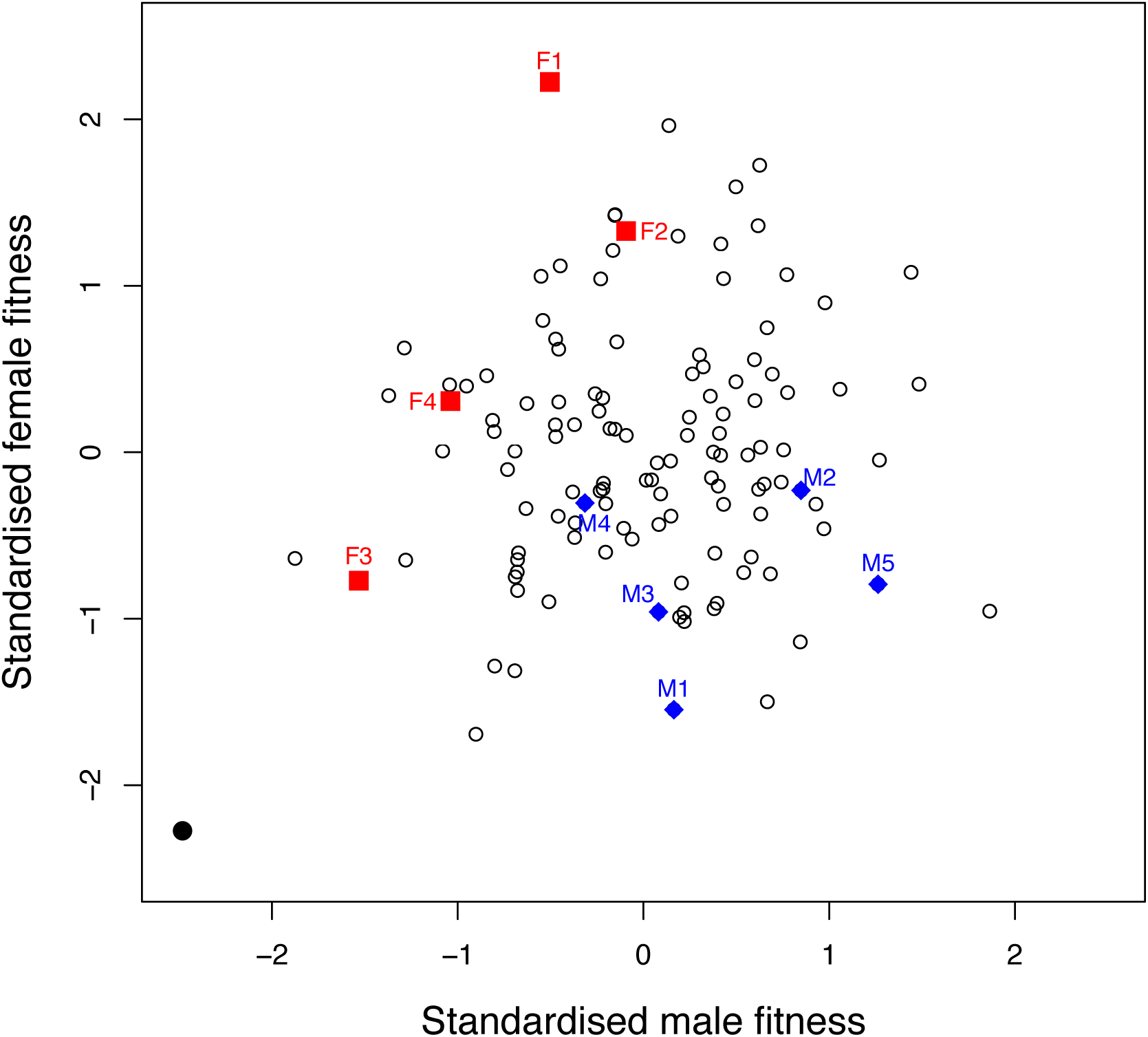
Male and female adult fitness across genotypes in the LH_M_-UCL population. Average male and female fitness across 113 hemiclonal lines randomly extracted from LH_M_-UCL (open circles). One line (filled black circle) showed extremely low fitness in both sexes and was removed from further analyses. Fitness measures obtained at UCL for a set of hemiclones from LH_M_-UU previously assayed as part of Innocenti and Morrow (2010) are also shown. The blue diamonds and red squares show the UCL fitness estimates of the hemiclones from this set that were classed as male beneficial/female detrimental and female beneficial/male detrimental fitness lines, respectively, in that previous study. Labels identify individual hemiclones for comparison with their fitness values in the previous study, shown in Figure S1.

Fitness assays for LH_M_-UCL were conducted in groups of 60 flies per vial, including ten focal flies, 20 standard *bw* competitors of the same sex and 30 standard *bw* flies of the opposite sex. For male assays, the flies were allowed to interact for 66 hours (days 11-14, interaction and oviposition phases of the 14-day rearing cycle) and fitness was measured as the proportion of offspring produced by the females of an assay vial that were sired by the focal hemiclonal males. For female assays, flies were allowed to interact for the 48 hours of the competition phase of the rearing regime (days 11 and 12 of the 14-day cycle) and fitness was measured as the average number of eggs laid by the focal hemiclonal females over the following 19.5 hours (the oviposition phase of the rearing regime). The average fecundity of *bw* competitors was also measured and included in standardised fitness measures (see section ‘Transformation of fitness data’).

**Fitness measurements for the LH_M_-UU population**: The dataset for the LH_M_-UU population had been compiled as part of Innocenti and Morrow (2010) and comprised fitness measures obtained from 100 hemiclones extracted from LH_M_-UU in October 2007. Fitness data had been obtained in a similar manner to that described above for the UCL population. Small differences included that fitness trials were performed on thirty flies per competition vial (five target individuals in competition with 10 *bw* flies) and that flies in the male assay were allowed to interact for 48 hours (instead of 48+18) before females were isolated (more details in Innocenti and Morrow 2010). Six male assays and four female assays were performed in the LH_M_-UU population, thus testing fitness of a total of 30 individual males and 20 individual females per hemiclone (compared to 30 of each in the LH_m_-UCL dataset).

### Statistical analysis of fitness

**Transformation of fitness data**: Before analysis, each set of fitnesses was standardised to remove block and vial effects. Thus, we defined fitness as the residuals of a linear model that decomposed raw individual fitness scores into a fixed effect of experimental block and random residual error (which includes the genotypic effect on fitness). For female fitness measures obtained for LH_M_-UCL, we further included a fixed effect describing the productivity of individual competition vials, as measured by the average number of offspring produced by *bw* competitor females. The residuals of these models were Z-transformed separately for each sex and population in order to obtain metrics of fitness that were comparable across sexes and populations.

**Comparing fitness measures across laboratories**: We used nine hemiclones from the LH_M_-UU sample to verify that fitness across populations was measured in a repeatable and comparable way. These hemiclones constitute the extremes of the fitness distribution of the LH_M_-UU sample (4 with extremely female-beneficial/male detrimental and 5 with extremely male-beneficial/female-detrimental effects) and had been maintained in the Uppsala laboratory. Complementing their fitness measures originally obtained in Uppsala, the fitness of these hemiclones was assayed again at UCL alongside those of the LH_M_-UCL sample, using exactly the same experimental protocol as that used for all other UCL hemiclones. To compare the fitness scores across laboratories, we performed correlation analyses, separately for each sex, between the standardised fitness scores obtained in the original analysis by Innocenti and Morrow (2010) and those obtained in the experiments at UCL. In addition, we applied Analyses of Variance, again separately for each sex, to the standardised fitness data from both laboratories, modelling fitness as a function of hemiclone (G), laboratory (E) and their interaction (GxE).

**Estimation and comparison of genetic variance components**: We used mixed models to estimate the contribution of additive genetic effects of hemiclones to the variation in male and female fitness, and the covariance between these genetic effects on fitness in males and females. Prior to analysis, we removed one outlier hemiclone from the UCL dataset that had a very low male and female fitness (Fig. 2), compatible with the effects of a strongly deleterious mutation affecting both males and females. Removing this outlier was conservative because, if included, it artificially increased the estimates of heritabilities and the intersexual genetic correlation in the UCL population.

We used WOMBAT (Meyer 2007) to fit the following multivariate animal model:

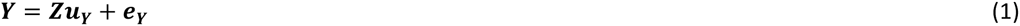

where ***Y*** is the vector of standardised fitness scores (hemiclones, sexes and populations concatenated), ***Z*** is the incidence matrix defining the population-specific combination of sex and genotype for each fitness value, ***u_Y_*** is the vector of sex- and population-specific additive genetic effects for fitness and ***e_Y_*** is the vector of residual effects (Meyer 1991; Lynch and Walsh 1998).

We estimated a cross-population genetic (co)variance matrix

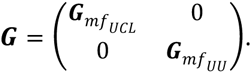

In this matrix, 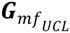and 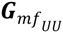are the population-specific additive genetic variance-covariance matrices,

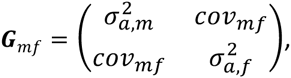
where 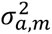and 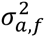are the additive genetic variances for fitness in males and females and *cov_mf_* is the inter-sexual additive genetic covariance. Due to the standardisation of our fitness data to a mean fitness of 0 and a standard deviation in fitness of 1 within each sex and population, heritabilities are directly given by additive genetic variances, 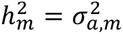and 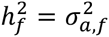. The inter-sexual genetic correlation for fitness can be calculated from the elements of the genetic variance-covariance matrix as 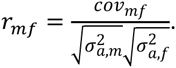

Estimation of genetic effects in the animal model relies on the numerator relationship matrix ***A*** that describes shared additive genetic effects between pairs of individuals. When defining this matrix, we considered hemiclonal males and females as full-sibs (*A_ij_* = 1/2 for individuals *i* and *j* that are part of the same hemiclonal line) and all other pairs as unrelated (*A_ij_* = 0 for pairs from different hemiclonal lines). The model was fitted using Restricted Maximum Likelihood (REML) and parameters and their approximate sampling errors (sensu Meyer and Houle 2013; Houle and Meyer 2015) were estimated.

The significance of parameter estimates and their differences between populations were tested using Log-likelihood Ratio Tests (LRTs) based on the X^2^ distribution. These compared the full model (eq. 1) in which all parameters were freely estimated to simpler nested models in which specific parameters had been fixed to appropriate values (function FIXVAR in WOMBAT). Specifically, we tested for within-population differences between the heritabilities 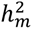and 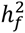of fitness by comparing the full model to a model where genetic variances 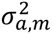and 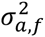were fixed to the average of their values estimated in the full model. To test whether the genetic covariance between male and female fitness *cov_mf_* within a population was significantly different from 0, we compared the full model to a model where the covariance term for that population was fixed to 0. Similarly, to test for between-population differences in either the male or female fitness heritability or the covariance between male and female fitness, we compared the full model to models in which the values of these parameters in each population were fixed to their average between the UU and UCL estimates obtained in the full model.

### DNA extraction and sequencing

We sampled 165 female adult flies from each of the populations in March 2012, about 260 generations after the separation of the two populations and about 46 and 115 generations after hemiclonal genomes had been sampled from LH_M_-UCL and LH_M_-UU, respectively. We extracted total genomic DNA from homogenised flies pooled by population using DNeasy Blood and Tissue Kit (Qiagen) and purified it using Agencourt AMPure XP beads (Beckman Coulter). One paired-end Illumina library (insert size <500bp) was made from each pool using the Nextera DNA Sample Preparation Kit (Illumina). Libraries were sequenced on an Illumina HiSeq2000 machine at the Centre for Genomic Research, University of Liverpool. Sequencing reads were extracted using CASAVA (version 1.82). Paired reads were trimmed using Sickle (version 1.2, default settings) and de-duplicated using Picard (version 1.77, http://picard.sourceforge.net) before being aligned to the *D. melanogaster* reference genome (BDGP5.25.60) using Bowtie2 (version2.0.0-beta7, option -X 500). We removed regions flanking indels (+/-5bp) with Popoolation (version 1.2.2, Kofler et al. 2011a) and used RepeatMasker (Smit et al. 2013–2015, version 4.0.5 with option species=drosophila and flag no_is) to mask interspersed repeats and low complexity regions. Finally, we applied a minimum read depth filter of 100 to ensure adequate precision of estimated allele frequencies, and a maximum read depth filter of 290 (about twice the average read depth of our sequencing runs, see Tab. S2), to avoid false positive SNPs due to duplicated genomic regions.

### SNP detection and analysis

We called SNPs in each population separately using SNVer (Wei et al. 2011, version 0.4.1 release 4, function for pooled sequencing SNVerPool.jar with minimum read and mapping quality cutoffs mq=20 and bq=20, haploid pool size n=330 and no filtering by minor allele frequency t=0). SNVer detects the significant presence of reads with the alternative allele (rather than polymorphism), so we re-ran the program on sites with high frequencies of the alternative allele, but using a reference genome sequence in which the corresponding positions were flipped to the alternative allele (i.e., testing for significant presence of the reference allele). We considered that a SNP was present if both polymorphism tests were significant. All P-values were corrected for false discovery rate (FDR<0.05) using the package qvalue (http://bioconductor.org). For all SNPs identified in this way, allele frequencies were extracted from the SNVer output. Furthermore, we calculated F_ST_ as a measure of genetic differentiation between LH_M_-UU and LH_M_-UCL using PoPoolation2 (Kofler et al. 2011b).

In order to summarise the chromosomal distribution of candidate SNPs with significantly elevated F_ST_ (see Results), we calculated the median distance between all pairs of adjacent candidate SNPs, separately for the X chromosome and the autosomes. We tested for significant clustering of candidate SNP by randomly permuting ‘candidate’ and ‘non-candidate’ labels among SNP loci and re-calculating the median distance among SNPs labelled ‘candidate’. Null distributions for the median distances between candidate SNPs were generated from 1000 such permutations (again, separately for X-linked and autosomal markers). A P-value for significant clustering was calculated as the proportion of median distances in the null distribution that was smaller or equal to the observed median distance.

### Functional characterisation of selected SNPs

We used the Variant Effect Predictor tool from Ensembl (McLaren et al. 2010) to map all SNPs to annotated genes and infer the consequences of variants (such as synonymous or non-synonymous coding sequence changes, splice variants, etc.). The tool uses extended gene regions that span 5 Kb up- and down-stream of the gene coordinates. In line with this default setting, we considered genes with candidate SNPs in that range as candidate genes. Analyses ignoring up-and downstream variants in these extended regions provided qualitatively identical results.

To assess the overlap between candidate genes in our study and previously described genes with sexually antagonistic expression patterns, we matched the identifiers of candidate genes to the corresponding Affymetrix *Drosophila* 2 probeset IDs. We then used the FlyAtlas data (Chintapalli et al. 2007) to consider only probesets that were expressed in adults, conservatively defined as those detected as ‘present’ in at least one library of one FlyAtlas adult tissue sample. The list of adult-expressed probesets covered by our study was then matched to the list of genes with antagonistic expression from Innocenti and Morrow (2010, supplementary table S1), retaining only those probesets in their study that were covered by our SNP data, and matches were back-translated into FlyBase gene identifiers.

We used Gene Ontology (GO) analysis implemented in DAVID (Huang et al. 2009a, b) to obtain insights into the functions of candidate genes. For enrichment analyses, we used all genes covered in our SNP dataset as the background set and applied a Benjamini-Hochberg correction for multiple testing to the P-values of individual tests. We further applied DAVID’s Functional Annotation Clustering (using the ‘high’ stringency setting) to categorise enriched individual GO terms into groups of related annotations.

## Results

### Hemiclonal fitness can be measured reliably across laboratories

A meaningful comparison between the genetic architecture of fitness across populations requires that the fitness of different genotypes be measured in a reliable and comparable way and in the absence of significant genotype-by-environment effects. Analyses of the fitness data from our nine reference hemiclones suggest that this is true here. First, we observed significant positive correlations between the sex-specific fitness measures obtained in the original experiments at the University of Uppsala and those obtained on the same hemiclones at UCL (females: r=0.81, t_7_=3.67, P=0.008; males: r=0.73, t_7_=2.79, P=0.027, Fig. S2). Second, ANOVAs performed on male and female fitness returned highly significant differences between hemiclones across laboratories, but no hemiclone-by-laboratory effects (females: hemiclone - F_8,45_=11.42, P<0.0001, hemiclone-bys-laboratory - F_8,45_=1.19, P=0.33; males: hemiclone - F_8,63_=11.34, P<0.0001, hemiclone-by-laboratory - F_8,45_=1.54, P=0.16; laboratory term non-significant in both analyses, as expected for Z-transformed fitness data). Based on the (additive) Sums of Squares, these analyses also show that, in line with the non-significant effect, the total variance in the data attributable to hemiclone-by-laboratory interactions is low (females: SS_hemiclone-by-laboratory_/SS_total_=0.065; males:SS_hemiclone-by-laboratory_/SS_total_=0.073; see Tab. S1 for full ANOVA tables). These results indicate that the fitness variation between genotypes detected in our experiments is overwhelmingly due to genetic differences, rather than the interaction between genotypes and the specific assay and laboratory environments.

### The intersexual genetic correlation for fitness differs between populations

We evaluated whether the genetic architecture of fitness, and in particular the inter-sexual genetic correlation for fitness, differed between the two replicate LH_M_ populations. Fitness heritabilities for LH_M_-UU were slightly higher than the estimates obtained in the earlier analysis of the UU dataset (Tab. 1 in Innocenti and Morrow 2010), in line with the fact that our approach accounted for the environmental effects of experimental assays. Our estimates confirmed that the heritability of male and female fitness differed in the LH_M_-UU population (X^2^=12.2, P=0.0005, Tab.1) while showing that LH_M_-UCL male and female heritabilities did not differ significantly (X^2^=0.7, P=0.39). Furthermore, male and female fitness heritabilities did not significantly differ across populations (X^2^=2.6, P=0.11 and X^2^=0.4, P=0.52, respectively).

While both populations featured ample and comparable heritable fitness variance, they differed in their inter-sexual genetic correlation. The point estimate of 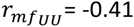was significantly negative (X^2^=4.5, P=0.03) while the genetic correlation in LH_M_-UCL was positive 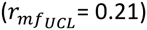and not significantly different from 0 (X^2^=1.2, P=0.27). Further, the intersexual genetic correlations differed significantly between populations (X^2^=5.1, P=0.02, Tab. 1, Figs. 2 and S1). This indicates that the sexual antagonism that was present in LH_M_-UU was absent in LH_M_-UCL.

### Population genomics revealed regions of significant genetic divergence between the populations

We performed genome-wide pooled sequencing of flies from LH_M_-UU and LH_M_-UCL in order to identify SNP loci with significant allele frequency differences between the populations (see Tab. S2 for general sequencing statistics). These loci would be candidates for regions potentially functionally related to the change in the genetics of fitness observed in the quantitative genetic analysis. Our sequencing approach covered the entire genome and for completeness we present results for all chromosomes, even though the contribution of the small 4th chromosome to fitness variation was not measured in our phenotypic assays.

Population sequencing identified more than 680,000 high-quality SNPs with significant allelic variation in at least one of the two populations. The density of SNP loci varied between chromosome arms (X_5_^2^=44997.1, P<0.0001). Chromosome arms 2L, 2R and 3L were enriched for SNP polymorphism, chromosome arm 3R had slightly fewer SNPs than expected, and chromosomes X and 4 were severely depleted for polymorphic sites (Tab. S3). The lower SNP densities on chromosomes X and 4 are expected based on the lower effective population sizes of these chromosomes, caused by the lower numerical population size of the X chromosome relative to the autosomes (Charlesworth et al. 1987; Mank et al. 2010) and selective interference along the virtually non-recombining chromosome 4 (Jensen et al. 2002; Haddrill et al. 2007; Betancourt et al. 2009; Charlesworth et al. 2010).

Analysis of the SNP allele frequencies showed that LH_M_-UU was weakly, but significantly, more genetically diverse than LH_M_-UCL. This difference was reflected at two levels. First, the percentage of SNP loci that were variable in LH_M_-UU but fixed in LH_M_-UCL (12.3%; Tab. S3) was greater than the percentage that only segregated in LH_M_-UCL (8.8%; Proportion test: X_1_^2^ = 4575.2, P<0.0001). Second, expected heterozygosity (H_e_) was slightly higher in LH_M_-UU than LH_M_-UCL (Fig. S3), both for autosomal SNPs (H_e-auto,UU_ = 0.295±0.165 mean±SD; H_e-auto,UCL_ = 0.278±0.173; paired t-test: t_613527_ = 73.3 78, P<0.0001) and for X-linked SNPs (H_e-X,UU_ = 0.284 ±0.166; H_e-X,UCL_ = 0.264 ±0.176; paired t-test: t_69303_ = 26.3 50, P<0.0001; note that these results are robust to corrections for possible non-independence between sites, e.g., including only every 10th, 20th or 50th SNP). These patterns also show again that X-linked variation was smaller than autosomal variation in both populations.

We estimated the genetic differentiation between the populations at each SNP locus by calculating the fixation index F_ST_ (Fig. 3 for chromosome arm 2L as an example, Fig. S4 for all chromosomes). The average level of differentiation across autosomal SNP loci was F_ST_ = 0.054±0.073, and that on the X chromosome was F_ST_ = 0.071±0.099. In order to identify SNP loci where genetic differentiation significantly exceeded the level expected from random genetic drift, we used the 105,851 SNPs causing synonymous variation (amino acid and stop codons). Synonymous allelic variation can be considered nearly neutral and used to establish an empirical null distribution of neutral background differentiation, against which sites under potential selection can be compared (e.g., Mcdonald and Kreitman 1991; Smith and Eyre-Walker 2002; Andolfatto 2005; Haddrill et al. 2010). We defined cut-off values for selective divergence between populations as F_ST_ values that exceeded the 99th quantile of the F_ST_ distribution across synonymous sites (Fig. S3). We did so separately for the X chromosome and the autosomes, in order to accommodate the different intensities of genetic drift acting on these two genomic compartments. The cut-offs used were F_ST_>0.371 for autosomes and F_ST_>0.453 for the X chromosome. Applying these cut-offs to our entire SNP dataset (excluding the synonymous SNPs, which were used to define the cut-offs), 3,762 autosomal and 717 X-linked SNP loci showed above-threshold levels of population differentiation (Fig. S5). The average absolute frequency difference between the two populations at these loci was 0.666±0.091 (median absolute difference 0.658). Independent verification using allele counts confirmed our F_ST_-based definition of population differentiation, as allele frequencies differed significantly between the populations at all candidate loci, even when using a stringent Bonferroni correction (Fisher’s Exact test on counts of reference and alternative alleles, P<0.05/4479=1.12 10^−5^ for all loci).

**Figure 3.**
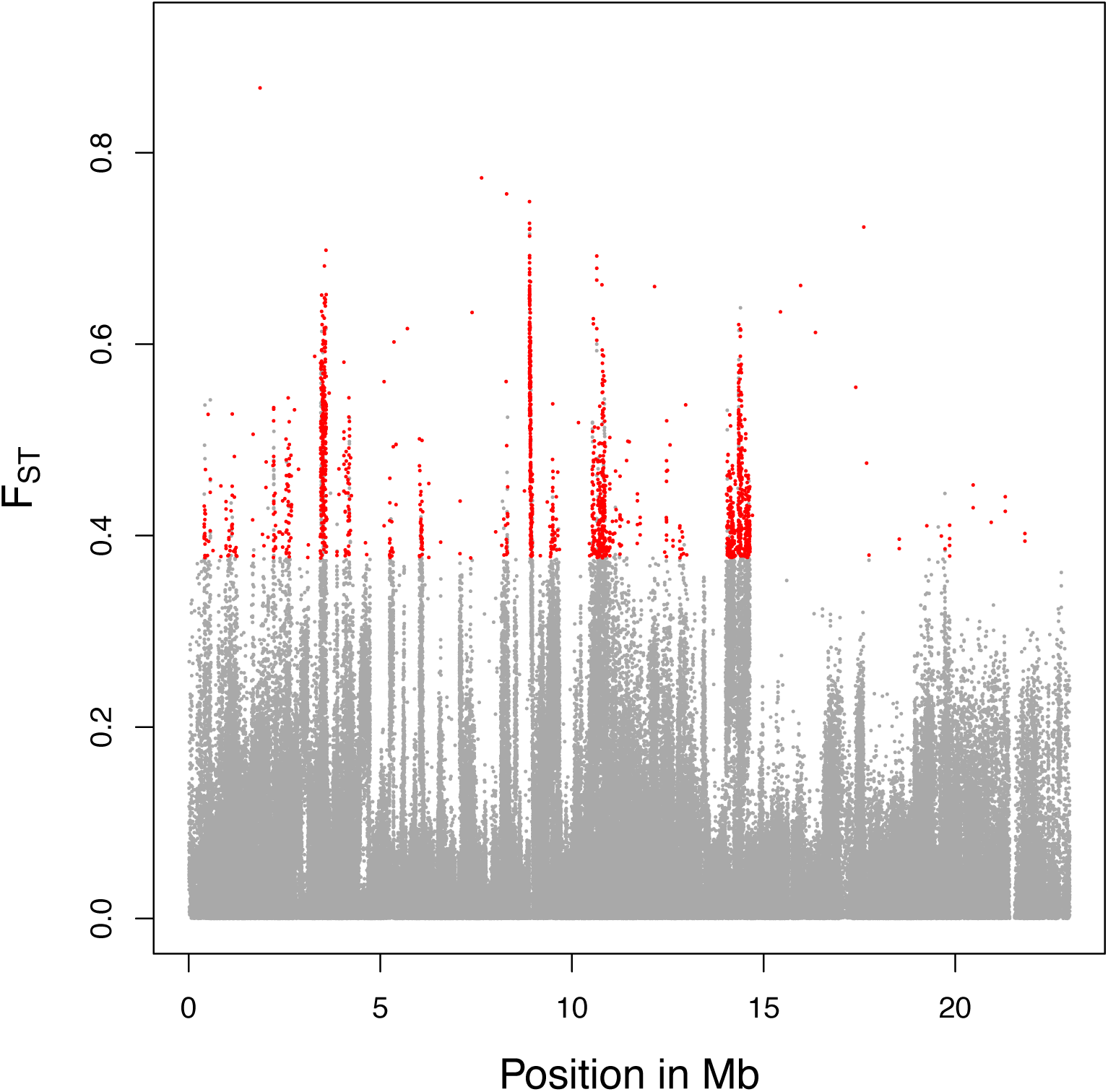
F_ST_ variations along SNPs in chromosome arm 2L. Grey dots indicate non-candidate loci, red dots candidate loci.

The distribution of the candidate SNPs was non-random across the chromosome arms and differed from that of non-candidate SNPs (Χ_5_^2^= 1082.1, P<0.0001). Specifically, candidate SNPs were over-represented on the X chromosome and chromosome arm 3R and under-represented elsewhere (Tab. 2). In addition to showing uneven distributions between chromosomes, candidate loci showed a clustered distribution along the chromosome arms and a large proportion of them fell within 100bp from each other (Fig. S6). A permutation test (see Methods) confirmed statistically significant clustering and showed that the observed distance between candidate SNPs was much smaller than expected by chance (autosomes: observed median distance = 770bp, range of median distances among 1000 sets of permuted loci = [12,998bp-14,978bp], P<0.001; X chromosome: observed median distance = 1,292bp, range of permuted median distances = [15,592bp-20,567bp], P<0.001).

The local clustering of candidate SNPs could indicate population differentiation in the frequency of chromosomal inversions, leading to parallel frequency changes of large numbers of alleles linked within the inverted part of the chromosome. Such effects have been observed in frequency clines among populations (Fabian et al. 2012) and frequency changes in response to laboratory selection (Kapun et al. 2014). In order to assess this possibility, we inspected patterns of polymorphism for diagnostic SNP alleles linked to seven cosmopolitan *D. melanogaster* inversions identified by Kapun et al. (2014). All but a few of these marker positions were well covered with high-quality reads in both samples (Tab. S4), but the data indicate that none of the inversions segregate in our populations. Only two of the inversion markers showed significant polymorphism in our samples (Tab. S4) and in both these cases the marker allele is the reference allele, suggesting either homoplasy (where the site on the inversion has convergently mutated back to the ancestral state) or an error in the inference of the marker allele.

### Candidate SNPs show biased patterns of allele frequency change

Population differentiation and elevated F_ST_ at candidate sites could arise due to allele frequency change in LH_m_-UCL, in LH_m_-UU or in both. In order to distinguish between these possibilities and infer the directionality of evolution at candidate SNPs, we used genotype data from the Drosophila Genetics Reference Panel (DGRP, Mackay et al. 2012; Huang et al. 2014). The DGRP constitutes an independent sample of genetic diversity from a wild North American population of *D. melanogaster* and— in the absence of data on the ancestral LH_M_ population—provides a suitable outside reference point. Focusing on the 636,924 SNPs that were shared and bi-allelic across the DGRP and LH_M_ samples, we found that levels of genetic differentiation between each of the LH_m_ populations and the DGRP differed markedly between non-candidate and candidate SNPs. For non-candidate SNPs, F_ST_ values between LH_M_-UCL and the DGRP were of similar magnitude—although marginally (and significantly) higher— than those between LH_M_-UU and the DGRP (autosomes: F_ST, UCL-DGRP_=0.099±0.127 mean±SD, F_ST,UU-DGRP_=0.091±0.118, mean pairwise difference=0.0073, CI=[0.0070,0.0076], t_568070_ =47.73 9, P<0.0001; X chromosome: F_ST,UCL-DGRP_=0.105±0.126, F_ST,UU-DGRP_=0.092±0.114, mean pairwise difference=0.0128, CI =[0.0118,0.0138], t_64464_=24.904, P<0.0001). For candidate SNPs, in contrast, we observed a large excess of differentiation between LH_M_-UCL and the DGRP, compared to that between LH_M_-UU and the DGRP (autosomes: F_ST,UCL-DGRP_=0.270±0.224, F_ST,UU-DGRP_=0.145±0.167, mean pairwise difference=0.124, CI=[0.113,0.136], t_3634_ =21.348, P<0.0001; X chromosome: F_ST,UCL-DGRP_=0.315±0.235, F_ST,UU-DGRP_=0.142±0.154, mean pairwise difference=0.173, CI =[0.146,0.201], t_707_=12.548, P<0.0001). Echoing these quantitative differences, a disproportionally large number of candidate loci showed greater F_ST_ between LH_M_-UCL and the DGRP than between LH_M_-UU and the DGRP (autosomes: 2327 of 3635 loci with F_ST,UCL-DGRP_> F_ST,UU-DGRP_, X_1_^2^= 202.660, P<0.0001; X chromosome: 458 of 708 loci, X_1_^2^= 34.168, P<0.0001). These results show that differentiation between the two LH_M_ populations at candidate sites is disproportionally driven by allele frequency change in LH_M_-UCL.

### Candidate SNPs have non-random functions

To understand the functional relevance of the candidate SNPs, we used the Variant Effect Predictor tool to examine the genetic consequences of variants segregating at candidate and non-candidate sites. When compared to non-candidate SNPs, candidate SNPs were over-represented among non-synonymous coding polymorphisms (amino acid changes, loss and addition of translation start sites, stop codons; Tab. 3). In contrast, we observed an under-representation of candidate SNPs in intergenic regions (i.e., those further than 5Kb away from any annotated gene). These non-random patterns of enrichment and depletion are indicative of a functional role for candidate polymorphisms.

Candidate SNPs mapped to a total of 1,131 genes, 939 on the autosomes and 192 on the X chromosome. We performed GO term enrichment analyses to gain information the biological processes in which these genes are involved and their molecular function. Term-by-term analyses revealed a strong association of candidate genes with biological processes related to growth, development and differentiation (Tab.S5A). These trends were further highlighted in subsequent clustering of enriched GO terms, where eight out of the ten most enriched term groupings were related to development (Tab. S6). In terms of molecular function, we found significant enrichment for two terms, both related to transcription regulation (Tab. S5B; no clustering was performed due to the small number of significant terms).

To relate our results to previous analyses of sexual antagonism, we compared our list of candidate genes to genes previously shown to have sexually antagonistic expression patterns. Innocenti and Morrow (2010) used a combination of phenotypic fitness assays and microarray expression analysis to identify genes that showed sex-differences in the relationship between expression level and fitness, mostly due to opposing associations of expression levels with male and female fitness (positive correlation between expression level and female fitness but negative correlation between expression level and male fitness, or vice versa). We observed a 30% excess of overlap between genes with candidate SNPs in our study and genes with such antagonistic expression patterns (147 overlapping genes to 112 expected, Χ_1_^2^=13.396, P=0.0003). No excess overlap was seen between our candidates and Innocenti and Morrow’s lists of genes showing expression levels that were only associated with male fitness (Χ_1_^2^=0.515, P=0.473) or only with female fitness (Χ_1_^2^=1.821, P=0.177). Thus, genes with antagonistic expression patterns were more likely to show large population differentiation in our study, but this association was not due to fitness-related genes being generally enriched among candidates in our study.

Finally, we investigated patterns of sex-biased gene expression among our candidate genes. Sex-biased expression is thought to be involved in the resolution of sexual antagonism. Accordingly, genes that show sex-biased expression should be less prone to antagonistic fitness effects and, inversely, genes with sexually antagonistic variation should have lower than average sex-biased expression. We compared our dataset to the Sebida database (Gnad and Parsch 2006) which classifies *D. melanogaster* genes as male-, female- or unbiased, based on the integration of multiple expression datasets. Our candidate genes show an under-representation of sex-biased gene expression, relative to non-candidate genes. This is true when pooling male- and female-biased genes (X_1_^2^=22.241, P<0.0001), and also when analysing male- and female-biased genes separately (male-biased: X_1_^2^=6.397, P=0.0114; female-biased: X_1_^2^=27.627, P<0.0001). This suggests that candidate genes tend to be those where, if present, antagonism could be expected to be more pronounced, as it is not tempered by sex-biased expression.

## Discussion

Despite the well-documented prevalence of sexual antagonism in plant and animal populations (e.g., Kohorn 1994; Chippindale et al. 2001; Fedorka and Mousseau 2004; Scotti and Delph 2006; Brommer et al. 2007; Foerster et al. 2007; Delph et al. 2011; Lewis et al. 2011; Mokkonen et al. 2011; Berg and Maklakov 2012) we know relatively little about its evolutionary dynamics (Cox and Calsbeek 2009; Stewart et al. 2010; Dean et al. 2012; Pennell and Morrow 2013). In this study, we document differences in intersexual genetic correlations of fitness (r_MF_) between two populations recently derived from the outbred laboratory stock LH_M_ in which sexual antagonism had originally been documented. While we found a significantly negative genetic correlation between male and female fitness in LH_M_-UU, consistent with ongoing SA in this population, r_MF_ was not different from zero and antagonism absent in LH_M_-UCL. These differences in the genetic architecture of sex-specific fitness were associated with significant shifts in allele frequencies at SNPs across the genome. These frequency changes occurred more often than expected at loci linked to genes that had been previously and independently linked to SA and more strongly affected frequencies in LH_M_-UCL, the population with reduced SA. We argue that our data are suggestive of rapid evolution of r_MF_ and are, at the very least, compatible with recent partial resolution of SA in LH_M_-UCL.

An important first step to interpreting our data is to establish whether evolutionary change has in fact occurred between the two populations. While this is straightforward at the genetic level, where we directly demonstrate differences in allele frequencies between the populations, caution is required when interpreting the observed difference in the genetic fitness correlation r_MF_. Gene-by-environment interactions are known to cause differences between estimates of genetic (co-)variances for male and female fitness obtained under different environmental conditions (Delcourt et al. 2009; Punzalan et al. 2014). Such effects could generate a spurious difference in r_MF_ here, because fitness measures for LH_M_-UU and LH_M_-UCL were obtained under conditions that—despite our efforts at standardisation—cannot be assumed to have been completely identical. Importantly, however, our inclusion of a reference set of genotypes in experiments at both Uppsala and UCL allowed us to rule out substantive genotype-by-laboratory effects. Comparison of fitness values between laboratories showed that the measures obtained at UCL correspond closely to those originally measured in Uppsala. Furthermore, the proportion of variation in male and female fitness that could be attributed to genotype-by-laboratory interactions was very small and not statistically significant. It therefore appears implausible that the observed differences in r_MF_ arose due to genotype-by-laboratory effects alone and can instead be assumed to reflect at least some evolutionary divergence in the genetic basis of fitness.

Several factors could have driven the evolutionary divergence between the populations. An important distinction is between neutral selective forces. Changes at the phenotypic (r_MF_) and allele frequency level can arise as a consequence of genetic drift during founding events or in populations of small effective size. Random fixation of antagonistic alleles can reduce the extent of SA or even render it undetectable. At the same time, however, such fixation events would result in a loss of heritable genetic variation in sex-specific fitness and a genetic homogenisation at the sequence level. These hallmarks of genetic drift do not appear in LH_M_-UCL, where antagonism was absent. Although it is difficult to compare absolute levels of quantitative genetic variation between populations, heritable variation in male and female fitness was present in LH_M_-UCL and tended to be greater than in LH_M_-UU. Similarly, our molecular data are incompatible with the action of strong genetic drift in LH_M_-UCL. Although the average heterozygosity was slightly reduced in LH_M_-UCL compared to LH_M_-UU, both populations showed comparable levels of allelic variation. Most importantly, loci with particularly elevated frequency differences showed many aspects of non-randomness in direction (divergence relative to the DGRP), position (clustering) and function (variant effects, association with antagonistic genes) that are incompatible with the action of random genetic drift. It therefore seems likely that the divergence between LH_M_-UU and LH_M_-UCL constitutes adaptive responses to selection pressures in one or both populations.

The most likely candidates for selection in these well-established laboratory populations are (i) differences in the micro-environment in which the populations were maintained, (ii) on-going co-evolutionary dynamics between the sexes (inter-locus sexual conflict) occurring independently within each population, or (iii) differences in the response to selection to alleviate the gender load (Rice 1992) that is generated by the sex-specific deleterious effects of sexually antagonistic alleles segregating in the populations. Our results cannot unambiguously differentiate between these alternative adaptive scenarios but we can assess the plausibility of each selective force in the light of the data. Environmental adaptation appears the least plausible scenario. It is conceivable that different selective pressure in the two laboratories could drive genetic divergence and potentially shifts in r_MF_, especially when considering pathogens as part of the environment. Even though LH_M_ stocks are superficially healthy, they are not maintained in sterile conditions and can be expected to undergo selection due to viral and/or bacterial pathogens. Interestingly, the evolution of immune resistance and disease tolerance shows sex-specific effects, where mutations altering resistance and tolerance can do so in opposing ways in males and females (Vincent and Sharp 2014). However, once again, divergent selection pressures deriving from any part of the environment in Uppsala and UCL would be expected to generate genotype-by-laboratory variation in fitness among our reference genotypes. In contrast, we found a positive correlation in their fitness across laboratories which implies that environmental selection pressures in the two laboratories were aligned, with the same genotypes (and hence phenotypes) having high (or low) fitness in both places. If environmental selection pressures occurred, their effect would have to be slight and unlikely to drive the population differentiation observed over the short timespans considered here.

Co-evolutionary arms races between the sexes occur whenever males and females differ in their reproductive interests. The promiscuous mating system imposed by the LH_M_ maintenance regime is expected to generate antagonistic interactions between males and females (‘inter-locus sexual conflict’ or simply ‘sexual conflict’, Chapman et al. 2003; Arnqvist and Rowe 2005), often over mating rates (Pischedda and Rice 2012).

The evolutionary arms race driven by sexual conflict can generate strong selection pressures, driving rapid divergence between populations (Rice and Holland 1997; Gavrilets 2000; Martin and Hosken 2003; Debelle et al. 2014). However, how this evolution would affect antagonistic variation is currently unclear. Potential links between male-female coevolution and sexual antagonism have been proposed (Pennell and Morrow 2013), but they await theoretical and empirical investigation. While we cannot rule out sexual conflict as a driver of evolution, certainly our data do not support that link. Many of the molecular phenotypes involved in the male contribution to the arms race (such as accessory gland proteins) are encoded by genes that show male-specific expression while our candidate SNPs are preferentially associated with genes that show unbiased expression in both sexes.

A more plausible scenario is that LH_M_-UCL adapted to selection generated by sexual antagonism. Sexually antagonistic variation can be stably maintained in populations under divergent selection on male and female traits (Gavrilets and Rice 2006), but any new variant that relieves the deleterious effect of antagonistic alleles in one sex while maintaining the benefit in the other is selectively favoured (Rice 1984). The invasion of such variants will contribute to the resolution of antagonism and the evolution of further sexual dimorphism. Several aspects of our data are compatible with such change occurring in the LH_M_-UCL population. First, the evolutionary change that we observe at the phenotypic and the genetic levels establish independent links to antagonism. This is obviously true for the phenotypes, where we directly assess SA. However, the population genomic analysis, which is unbiased with regard to the phenotypic measures, also invokes SA through the significant overlap of our candidates and genes previously shown to have antagonistic expression patterns (Innocenti and Morrow 2010). Second, the directionality of phenotypic and genetic evolution indicate adaptation in LH_M_-UCL, rather than symmetric divergence between the populations. The presence of SA and a negative rMF is the ancestral state in the LHM population, demonstrated in the original stock population at UCSB (Chippindale et al. 2001) and then confirmed in LH_M_-UU (Innocenti and Morrow 2010). The absence of antagonism (r_MF_≈0) thus is a derived state towards which LH_M_-UCL evolved. The same directionality is reflected at the genomic level. Our analysis of allele frequencies across the two LH_M_ populations and the DGRP show that frequency change at candidate loci consists disproportionately of LH_M_-UCL frequencies shifting away from those in LH_M_-UU. Phenotypic and genetic data thus paint mirror images of rapid change occurring mostly in LH_M_-UCL.

Our evidence in favour of a possible recent alleviation of antagonism in LH_M_-UCL contrasts with the repeated detections of antagonistic variation in LH_M_ (Chippindale et al. 2001; Innocenti and Morrow 2010). This suggests that the recent rapid evolution of SA occurred following a long period of stasis, during which antagonistic variation was stably maintained for many generations. It implies that the alleviation of SA was caused by a key innovation decoupling male and female phenotypes, most likely relying on a rare mutational event, such as several epistatic variants arising more or less simultaneously. It seems implausible that the many loci across the genome at which we observe significant population divergence are causally involved in the alleviation of antagonism. Instead, the frequency change we observe at most loci is more likely a consequence, rather than the cause, of alleviation. An emergent mechanism of SA resolution would alter the sex-specific fitness effects of antagonistic alleles previously maintained in balanced polymorphism, leading to shifts in frequencies and potentially the fixation of the allele with the greater average fitness across the sexes (Rice 1984; Connallon and Clark 2012; Mullon et al. 2012).

Analyses to identify the causal change or changes underlying an alleviation of SA are beyond the scope of this study. However, given the proviso that most of our candidate genes are associated with antagonism, our results are informative about the genetic basis of SA. Our data suggests an enrichment of loci on the X chromosome, in line with previous theoretical predictions (Rice 1984). It further suggests that the disproportionate contribution of the X chromosome to quantitative genetic variation in fitness previously observed in LH_M_-UCSB (Gibson et al. 2002) is at least in part due to the number of X-linked antagonistic loci (rather than their phenotypic effects). Our data also provide a first glimpse of genes potentially involved in antagonism. At a general level, the prominent involvement in development of our candidate genes implies that many antagonistic fitness effects may be rooted in anatomical differences between the sexes. This fits with data showing how developmental processes allow the decoupling of male and female phenotypes (Gompel et al. 2005; Roberts et al. 2009; Khila et al. 2012). At the level of individual genes, our candidates include several genes known to be involved in male courtship behaviour *(cacophony, period* and *Btk family kinase at 29A)* and loci related to sex determination and differentiation *(transformer 2, Wnt oncogene analogue 2, doublesex-Mab related 93B, sister-of-Sex-letha* and *bric ά brac)*. While these manual screens are subjective, the presence of such genes raises the intriguing possibility that some of the adaptive trade-offs between males and females are caused by variation close to the top of the sex-specific regulatory cascade.

In conclusion, we have presented data indicating rapid evolutionary change in the genetic basis of fitness and consistent with a partial resolution of antagonism in LH_M_-UCL. Due to the enormous scale of the phenotypic assays that we report, our study only covered a very small number of populations and relied on combining existing and new data that were collected at different time-points. Despite these limitations, our results demonstrate parallel change at phenotypic and genetic levels, establishing independent links to sexual antagonism. In the future, improved knowledge of the genetic basis of antagonism will help to identify the constraints acting on male and female traits and elucidate the genomic innovations that have made it possible for LH_M_-UCL to overcome such constraints.

## Acknowledgements

We thank Chris Wheat and three anonymous reviewers, whose comments greatly improved that quality of our manuscript. We thank Jessica White, Rebecca Finlay, and Anja Schlott for technical assistance with data collection, Adam Chippindale for his input to planning the fitness assays and Steve Chenoweth, Julie Bertrand, Jacqueline Sztepanacz, and Jarrod Hadfield for their assistance with the quantitative genetic analyses. Steve Paterson and Ian Goodhead (Centre for Genomic Research, University of Liverpool) assisted with the initial treatment and quality-control of sequencing data. JMC and SF were funded by a NERC research grant (NE/G019452/1) to MR and KF, JH by a BBSRC PhD studentship and MSH by a UCL IMPACT PhD studentship. PI and EHM were funded by the Swedish Research Council (2004-2572, 2006-2848, 2008-5533), a European Research Council Grant (#280632) and a Royal Society University Research Fellowship.

**Table 1.** Heritabilities and genetic correlations for fitness in LH_M_-UU and LH_M_-UCL.

| Population | Female $h^2$ | Male $h^2$ | $r_{mf}$ |
| --- | --- | --- | --- |
| LH<sub>M</sub>-UU | 0.71 (0.15) | 0.19 (0.07) | -0.41 (0.18)<sup>a *</sup> |
| LH<sub>M</sub>-UCL | 0.58 (0.12) | 0.41 (0.12) | 0.21 (0.19)<sup>b</sup> |
| Difference between populations | $X_1^2 = 0.4$, $P = 0.52$ | $X_1^2 = 2.6$, $P = 0.11$ | $X_1^2 = 5.1$, $P = 0.02$ |

The table provides estimates and, in parentheses, the approximate sampling error for the heritabilities of male and female fitness and the intersexual genetic correlation ( $r_{mf}$ ) for fitness in the two populations, as well as the results of likelihood ratio tests comparing estimates between populations (see Methods).
<sup>a</sup> Likelihood ratio test comparing the estimate to zero (see Methods): $X_1^2 = 4.5$ , $P = 0.03$
<sup>b</sup> Likelihood ratio test comparing the estimate to zero (see Methods): $X_1^2 = 1.2$ , $P = 0.27$

**Table 2.**
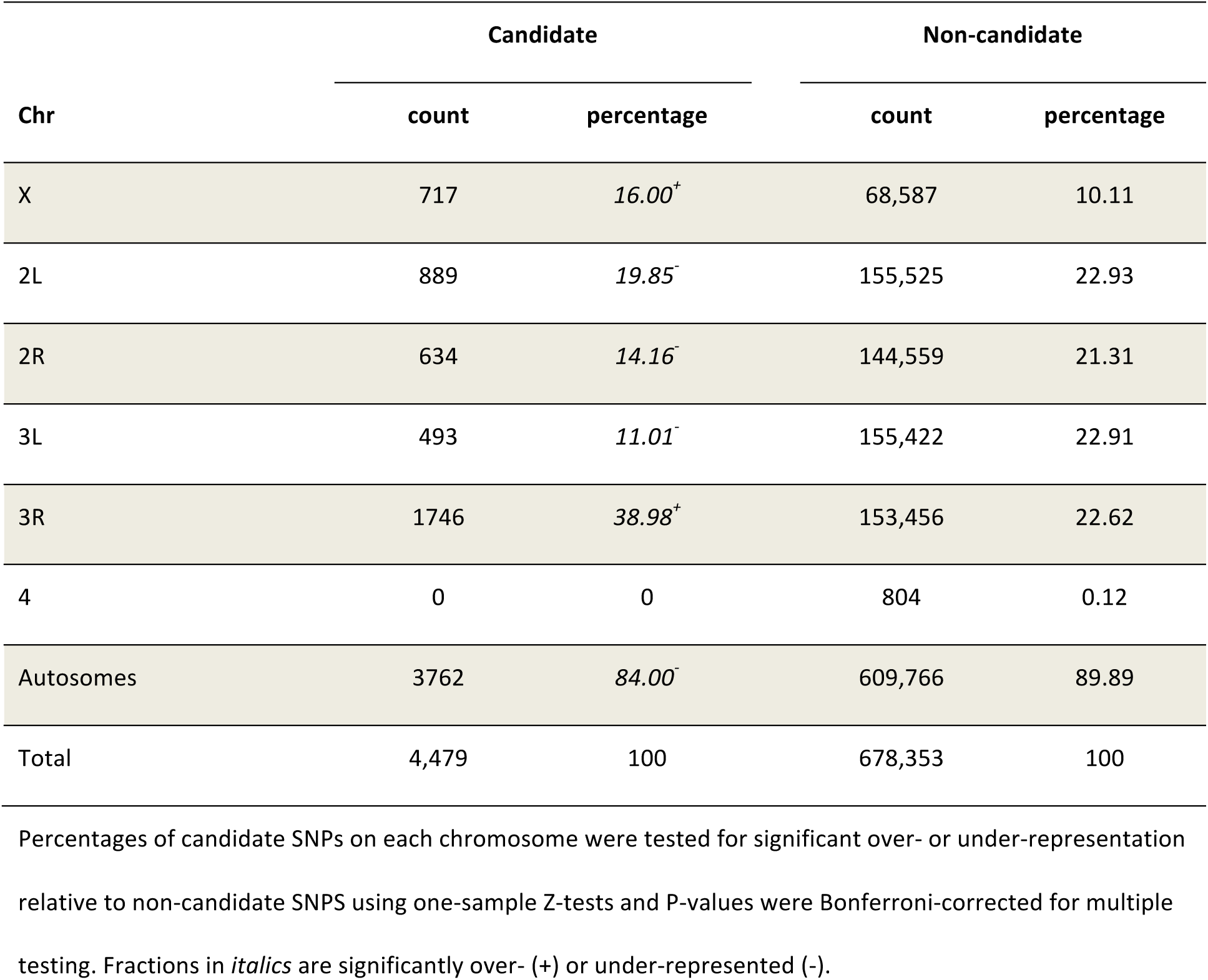
Distribution of candidate and non-candidate SNP loci across chromosomes.

**Table 3.** Comparison of genomic feature distributions between all SNPs, candidate SNPs and non-candidate SNPs.

| Functional category | All |  | Candidate |  | Non-candidate |  |
| --- | --- | --- | --- | --- | --- | --- |
|  | count | percentage | count | percentage | count | percentage |
| UTR | 42,706 | 6.25 | 312 | 6.97 | 42,394 | 7.41 |
| Non-synonymous | 33,483 | 4.90 | 312 | *6.97<sup>+</sup>* | 33,171 | 5.79 |
| Synonymous | 105,851 | 15.50 | 0 | – | 105,851 | – |
| Splice site | 6,484 | 0.95 | 61 | 1.36 | 6,423 | 1.12 |
| Intron/non-coding exon | 289,792 | 42.44 | 2,286 | 51.04 | 287,506 | 50.22 |
| Up-/downstream | 135,364 | 19.82 | 1,110 | 24.78 | 134,254 | 23.45 |
| Intergenic | 69,152 | 10.13 | 398 | *8.89<sup>-</sup>* | 68,754 | 12.01 |

Proportional representations of functional categories were tested for significant over- or under-representation of candidate SNPs compared to non-candidate SNPs using one-sample Z-tests and P-values were Bonferroni-corrected for multiple testing. Categories in *italics* are significantly over- (+) or under-represented (-).
Synonymous polymorphisms were not considered while performing these tests, as they had been used to identify candidate SNPs and hence not tested here. For easier comparison, the synonymous variants were also excluded when calculating the percentages of candidate and non-candidate variants that fall into the different categories.

